# The long non-coding RNA MALAT1 modulates NR4A1 expression through a downstream regulatory element in specific cancer-cell-types

**DOI:** 10.1101/2023.03.09.531856

**Authors:** Sara Wernig-Zorc, Uwe Schwartz, Paulina Martínez, Josefa Inalef, Francisca Pavicic, Pamela Ehrenfeld, Gernot Längst, Rodrigo Maldonado

## Abstract

Chromatin-associated long non-coding RNAs (lncRNAs) have been shown to define chromatin density, regulate gene expression, and are involved in the initiation and progression of various cancer types. Despite the wealth of studies describing transcriptome changes upon lncRNA modulation, little data is showing the direct effects of lncRNA on regulatory elements (REs) that drive gene expression. Here we explored the molecular mechanism of the chromatin-interacting lncRNA, MALAT1, through RNA- and ATAC-seq, using HeLa cells as a model system. Time-resolved MALAT1 knock-down assays revealed its direct regulation of a limited number of protein-coding genes. Loss of MALAT1 resulted in a substantial loss of chromatin accessibility downstream of the *NR4A1* gene, associated with its down-regulation. CRISPR-i assays revealed that this region corresponds to a new downstream RE. Next, using TCGA data, we identified a direct correlation between the expression of NR4A1 and the accessibility of the downstream RE in breast cancer. The molecular mechanism was validated on estrogen receptor (ER) positive breast cancer cells (MCF7) and Pancreatic Duct Epithelioid Carcinoma (PANC1) cells, not showing this effect according to TCGA data. Indeed, MALAT1 regulates the expression of NR4A1 in a cell type-specific manner by changing the accessibility of the downstream RE. MALAT1 exhibits a molecular mechanism that fine-tunes the expression of cancer drivers, like NR4A1, in ER-positive breast cancer cells, but not in other cell types.

## Introduction

Histones, chromatin-associated proteins, and RNA play essential roles in fine-tuning gene expression. Among them, chromatin-associated RNAs have arisen as key factors modulating nuclear architecture, genome-wide chromatin interactions, gene expression, and genomic integrity (1). Long non-coding RNAs (lncRNAs) are transcripts of at least 200 nucleotides in length, transcribed by RNA polymerase II, in most cases spliced, 3’ polyadenylated, and 5’ capped by 7-methyl guanosine (2). LncRNAs act as cis- or trans-acting transcriptional regulators depending on their functional interactions with chromatin, chromatin-associated factors, and nuclear ribonucleoprotein particles (3). Combining the information from different databases, almost 57.000 lncRNA genes have been described to generate more than 127.000 different transcripts from the human genome (4). Their expression is highly tissue and cell-type specific. The regulation of stem cell differentiation and renewal, in combination with specific protein partners, demonstrates their remarkable transcriptional regulatory potential (5). Lineage engagement for adipose, skeletal, cartilage, muscle, neuronal, and skin cell differentiation, in addition to embryonic stem cell pluripotency, have been described guided by different lncRNAs like PU.1 AS, Msx1as, HOTTIP, H19, HOTAIRM1, TINCR, RoR, Xist, Airn, and Firre, among others (54). Accordingly, it is not surprising that lncRNAs were linked with the onset and progression of cancer, regulating gene expression at transcriptional and post-transcriptional levels, and impacting the intermediary metabolism and cell signaling (2,6– 8).

The metastasis-associated lung adenocarcinoma transcript 1, MALAT1, is one of the most abundant lncRNAs in human cells, being located in nuclear speckles (9). In humans, MALAT1 is an 8 Kb single exon transcript expressed from chromosome 11q13 (10). Studies on the mouse homolog of the human *MALAT1* gene showed the RNaseP-RNaseZ post-transcriptional processing of MALAT1, to generate a long transcript retained on nuclear speckles and a 61 nt small RNA with a tRNA-like structure (11). Interestingly, after RNase processing, the 3’ end of the highly expressed long MALAT1 transcript, folds to generate a blunt-end triple helical structure that stabilizes the transcript through the protection from 3’-to-5’ exonucleases (12,13).

Different transcriptional-related processes are regulated by MALAT1. RNA-RNA interactions characterized by RNA Antisense Purification (RAP) sequencing experiments in mouse embryonic stem cells, showed that MALAT1 interacts mainly with pre-mRNAs. This interaction is mediated through proteins involved in alternative splicing, like the arginine/serine-rich phosphoproteins (SR proteins) (14). Accordingly, MALAT1 is predicted to regulate pre-mRNA splicing through the interaction with SR proteins, modulating their phosphorylation status, and influencing the distribution of other splicing factors in human cells (15). Comparing the results of RAP sequencing in mouse cells with data obtained by Capture Hybridization Analysis of RNA Targets (CHART) in MCF7 breast cancer cells, MALAT1 was found to interact also with DNA at numerous genomic sites. MALAT1 is preferentially associated with the coding region and termination or polyadenylation sites of transcriptionally active and spliced genes (14,16). Moreover, combining mass spectrometry with the CHART assay (CHART-MS), revealed the preferential interaction of MALAT1 with RNA processing factors, like nuclear speckle and paraspeckle proteins, and transcriptional regulators (16).

Different types of cancers have been associated with alterations of MALAT1 expression and/or function. MALAT1 was shown to act as a tumor suppressor or a tumor promoter, depending on the cancer type examined, or it can even have opposing roles in the same cancer type (6,17,18). The diversity of MALAT1 functions may be explained by its binding to regulatory genomic domains, its interaction with chromatin-modifying complexes, leading to the deposition of active or inactive histone marks, or its interaction with transcription factors that directly target gene regulatory elements (19–21). These data indicate that MALAT1 is associated with the regulation of gene expression during cancer development. Nevertheless, despite the diverse effects of lncRNAs (1), the molecular mechanism of MALAT1 function on gene expression in cancer is still unresolved.

In this study, we used HeLa cells as a model system to validate MALAT1 association with active chromatin. Then, modulating MALAT1 expression with Locked Nucleic Acid (LNA) Gapmers, we defined direct and indirect transcript targets by RNA-seq. These targets exhibited changes in chromatin accessibility of key regulatory elements as shown by ATAC-seq. Using the CRISPR-i technology, we found that MALAT1 modulates the accessibility of a downstream regulatory element of the nuclear receptor subfamily 4 Group A Member 1 (NR4A1). NR4A1 is a transcription factor dysregulated in different cancer types, involved in T-cell dysfunction, and associated with the development of fibrotic diseases (22–27). We correlated chromatin accessibility changes at the NR4A1-RE with NR4A1 gene expression using TCGA data. Interestingly, we observed a direct correlation for certain cancer types, such as breast cancer, but not for others. Furthermore, we confirmed this specific cancer-type association using CRIPR-i, showing that pancreatic ductal adenocarcinomas (PDAC), are not regulated by MALAT1 and do not change the expression of NR4A1. However, the NR4A1 gene in MCF7 breast cancer cells was specifically regulated by MALAT1. These results delineate the MALAT1 molecular mechanism that fine-tunes the expression of specific cancer modulators, like NR4A1.

## Results

### MALAT1 is a chromatin-associated lncRNA linked to the regulation of protein-coding genes

MALAT1 has been described as a chromatin-associated lncRNA, using different RNA-centered experimental approaches based on proximity ligation or antisense oligonucleotides (14,16,28–31). We performed a chromatin-centered method to identify lncRNAs associated with active chromatin. To isolate the active fraction of chromatin, the nuclei of crosslinked HeLa cells were isolated and the cytoplasmic RNA background was removed by several wash steps (Figure 1A). The active chromatin fraction was pulled down with antibodies directed against H3K27ac. As a negative control, the H3K27ac antibody was pre-bound with the acetylated H3 peptide to block epitope binding (Figure 1B). The chromatin-associated RNA was purified and analyzed by automated electrophoresis (Figure 1C). The input (RNAs from the nuclear extract) and negative control samples showed the same RNA distribution pattern, while the RNAs associated with H3K27ac marked chromatin revealed a different pattern, being highly enriched in RNAs of around 200 nucleotides (Figure 1C, Supplementary Figure 1A). RNAs were used for library preparation and sequencing (ChRIP-seq) (32,33). Quantification of the RNAs revealed that the H3K27ac-associated RNAs are highly enriched in promoter-derived protein-coding transcripts when compared to Input and the negative control (Figures 1D, 1E, Supplementary Table 1). Interestingly, about 5% of these transcripts correspond to lncRNAs (Figures 1D, Supplementary Table 1). We identified several lncRNAs that have been previously associated with chromatin, like MALAT1, NEAT1, Firre, and TARID (16, 54, 55) (Figures 1F and Supplementary Table 1). We further focused our study on MALAT1, being specifically enriched within H3K27Ac-associated transcripts, compared to input and negative control ChRIP data (Figure 1G). Next, the MALAT1-H3K27Ac interaction was validated by real-time PCR analysis of the ChRIP samples (Figure 1H). To identify direct and indirect transcriptional-target genes, we used LNA Gapmers to knock down (Kd) MALAT1 and control Gapmers to target LacZ. The MALAT1 Kd effects were assessed by RNA-seq and real-time PCR at 24- and 48-hours post-transfection (Figures 1I and 1J). These techniques validated the efficient MALAT1 Kd at both time points. Interestingly, after 24 hours, the MALAT1 Kd revealed a higher number of downregulated transcripts (313) compared to upregulated ones (143). Among them, the vast majority of differentially expressed transcripts correspond to protein-coding genes (Figures 1K, 1L, and Supplementary Table 2). After 48 hours, the number of differentially expressed transcripts strongly increases, reaching similar levels for up- and down-regulated transcripts (Figures 1K and 1L). At this time point, besides the preponderance of protein-coding genes, we started to observe the differential expression of non-coding genes (Figure 1L and Supplementary Tables 3-4). Gene ontology analysis by overrepresentation test (56, 57) of the 48 hours sample revealed that up-regulated genes are enriched in transcripts being associated with transcriptional regulation, compared to the 24-hour sample (Supplementary Figures 1B and 1C). The differences observed between the 24h and 48h time points suggest two important aspects of the function of this lncRNA. First, MALAT1 is primarily associated with the regulation of protein-coding genes. Second, the time-dependent changes of the effects suggest the presence of early direct targets (after 24h) and the emergence of indirect targets 48 hours after knock-down, when transcriptional regulators (proteins and nc-RNAs) show up.

**Figure 1.**
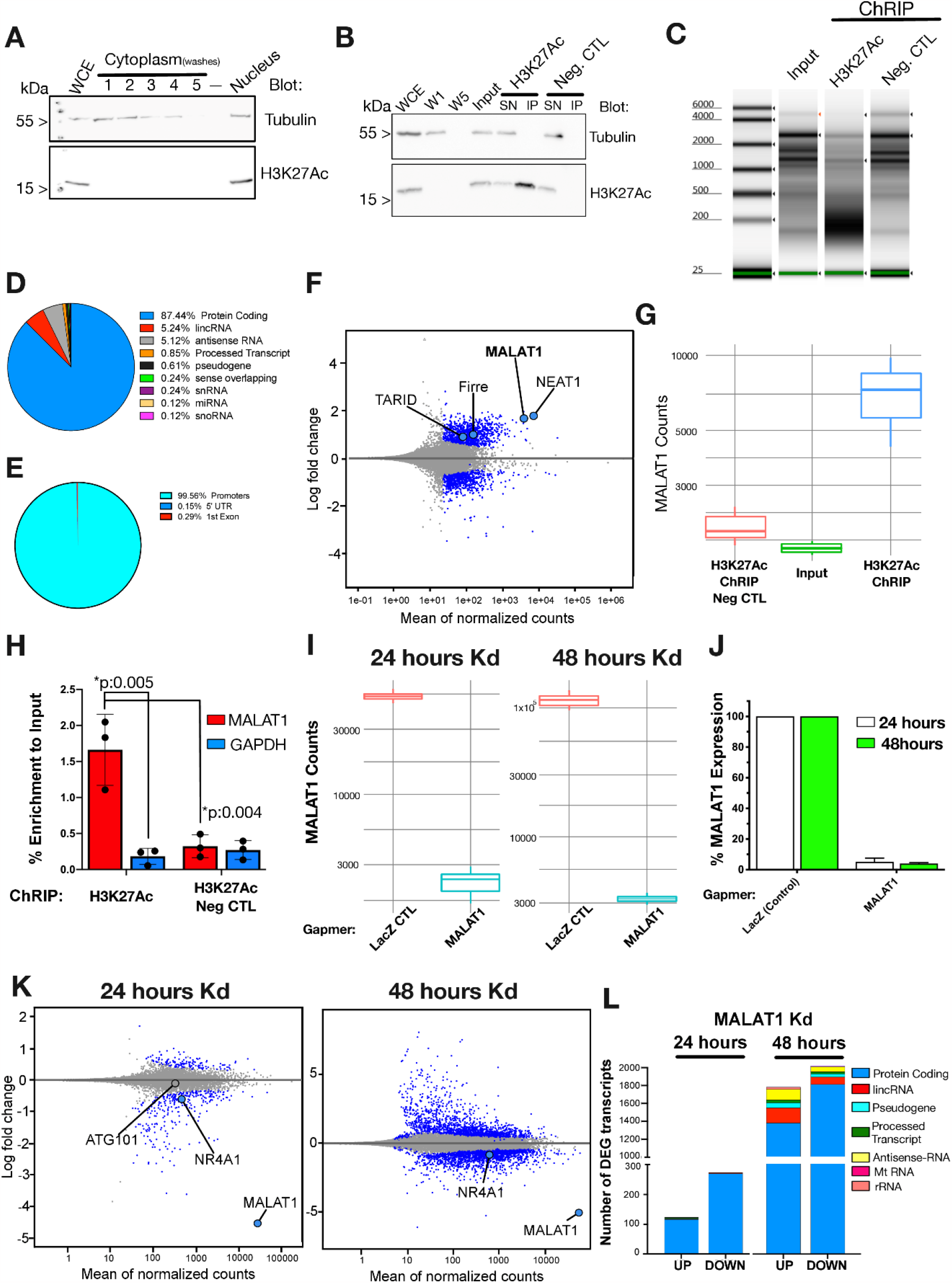
The chromatin-associated lncRNA MALAT1 modulates the expression of mainly protein-coding genes. **A)** Western blot analysis of HeLa cell fractionation using Tubulin and H3K27Ac as cytoplasmic and nuclear fractions markers, respectively. Whole-cell extract (WCE), cytoplasmic washes, and input fractions are shown. **B)** H3K27Ac immunoprecipitation verification by Western blot analyzing the WCE, the first (W1) and last (W5) cytoplasmic washes, the nuclear input, the unbound supernatant (SN), and immunoprecipitated (IP) fractions. **C)** H3K27Ac-derived RNAs automated-electrophoresis profiles. **D)** Gene type distribution of H3K27Ac-associated RNAs. **E)** Transcript peak annotation distribution of H3K27Ac-associated RNAs. **F)** MA-plot showing significantly enriched H3K27Ac-associated RNAs in blue. **G)** MALAT1 RNA-seq counts on each H3K27Ac ChRIP sample type. **H)** Real-time PCR detection of MALAT1 from ChRIP samples, GAPDH was amplified as an unspecific gene control. Significant differences are shown with p-values obtained from t-tests analyses. **I)** MALAT1 RNA-seq counts after 24 and 48 hours of MALAT1 Kd using specific Gapmers. LacZ Gapmers were used as a Kd negative control. **J)** Real-time PCR detection of MALAT1 after 24 and 48 hours of MALAT1 Kd. **K)** MA-plot for RNA-seq after 24 and 48 hours of MALAT1 Kd. Significant up and downregulated transcripts are indicated in blue. **L)** Quantification of significantly up and downregulated transcripts from RNA-seq data after 24 and 48 hours of MALAT1 Kd.

These results validate that MALAT1 is a chromatin-associated lncRNA bound to active chromatin, regulating the expression of specific protein-coding genes.

### MALAT1 modulates the chromatin accessibility of an *NR4A1* downstream regulatory element

To understand the effects after MALAT1 downregulation and to address the chromatin structure of the target genes, we performed ATAC-seq experiments 24 and 48 hours after MALAT1 Kd. By comparing the accessibility changes between both time points, we identified genomic loci that become inaccessible 24 hours after MALAT1 Kd and maintain this condition after 48 hours (Figure 2A, Cluster 4). Considering time-dependent changes in gene expression after MALAT1 downregulation, and the presence of putative direct MALAT1 targets after 24 hours of Kd, we focused on the early structure changes. Compared to the controls we identified numerous significantly altered ATAC sites with increased or decreased chromatin accessibility, distributed over all chromosomes (Supplementary Figure 2A). Nevertheless, these sites were not randomly distributed throughout the genome. Quantification of the ATAC-site distribution showed significant enrichments (Chromosome 1, 5, 6, 8, 11, 12, 15, 17, and 20) or reductions (Chromosome 4, 9, 13, 14, 16, 21, and X) of ATAC peak density (Figure 2B). Furthermore, the sites that change their accessibility already after 24 hours of MALAT1 knock-down were enriched in regulatory elements. Analyzing chromatin states, our data show that enhancers are the main regulatory elements affected after the MALAT1 downregulation. This was observed for both pools, ATAC sites that gained (Figure 2C) and lost (Figure 2D) accessibility after 24 hours of MALAT1 Kd. Analyzing the correlation between gene expression changes and changes in chromatin accessibility, revealed a higher overlap between chromatin closure and transcription de-regulation than the other way around (Figure 2E). This indicates that the sites that become less accessible after MALAT1 Kd have a stronger impact on gene expression compared to the sites that gain accessibility.

**Figure 2.**
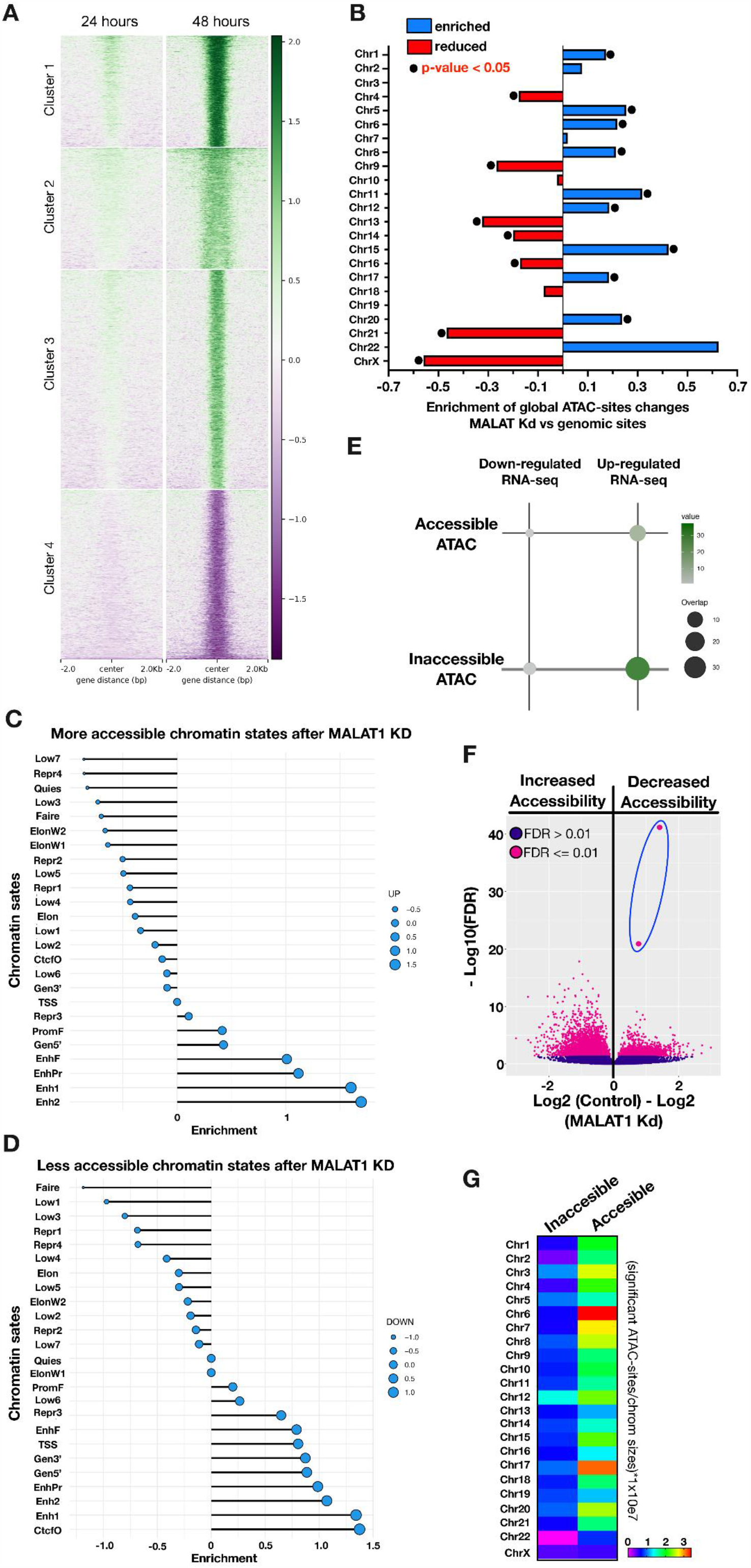
Chromatin accessibility changes after MALAT1 downregulation. **A)** Heatmap showing the K-means clusters of differentially accessible ATAC-seq sites of MALAT1 Kd versus control after 24 and 48 hours. Accessibility scores are shown on the right side. **B)** Enrichment of global ATAC-seq sites per chromosome ((%MALAT1 Kd sites/%genomic sites)-1). **C)** and **D)** Enrichment of significant ATAC sites that gained or lost accessibility after MALAT1 Kd 24 hours over chromatin states on the human genome. **E)** Overlap quantification of significant accessible/inaccessible ATAC-seq sites with differentially expressed genes after MALAT1 Kd. **F)** Quantification of significant ATAC-seq that changes their accessibility after MALAT1 Kd. **G)** Chromosomal distribution of significant accessible/inaccessible ATAC-seq sites after MALAT1 Kd.

We now focused on two such sites, exhibiting the largest decrease in accessibility after 24 hours of MALAT1 Kd (Figure 2F, dots inside the blue oval). These sites are part of cluster 4 (Figure 2A), and these are located on chromosome 12, a chromosome exhibiting a higher number of repressed sites than average (Figure 2G). The repressed ATAC-sites are located downstream of the *NR4A1* gene (Chr12: 52.022.832-52.059.507 on GRCh38), in a region flanked by additional strong ATAC-peaks and coinciding with CTCF-ChIP-seq peaks (red and black squares, respectively, Figure 3A). This region may relate to a topologically-associated domain (TAD), in which the *NR4A1* and *ATG101* genes are expressed as observed by our RNA-seq data (Figure 3A). Additionally, the active histone marks H3K27Ac/H3K4me3/H3K4me1 sign the *NR4A1* and *ATG101* promoters, and enhancer regions up and downstream of the genes (Figure 3B). Cap-Analysis of Gene Expression (CAGE) data on HeLa cells showed the location of the transcriptional start site, corresponding to the NR4A1 transcript ENST00000394825. The promoter and coding region of this transcript covers the whole ATAC-accessible region, including the downstream element (Supplementary Figure 2B).

**Figure 3.**
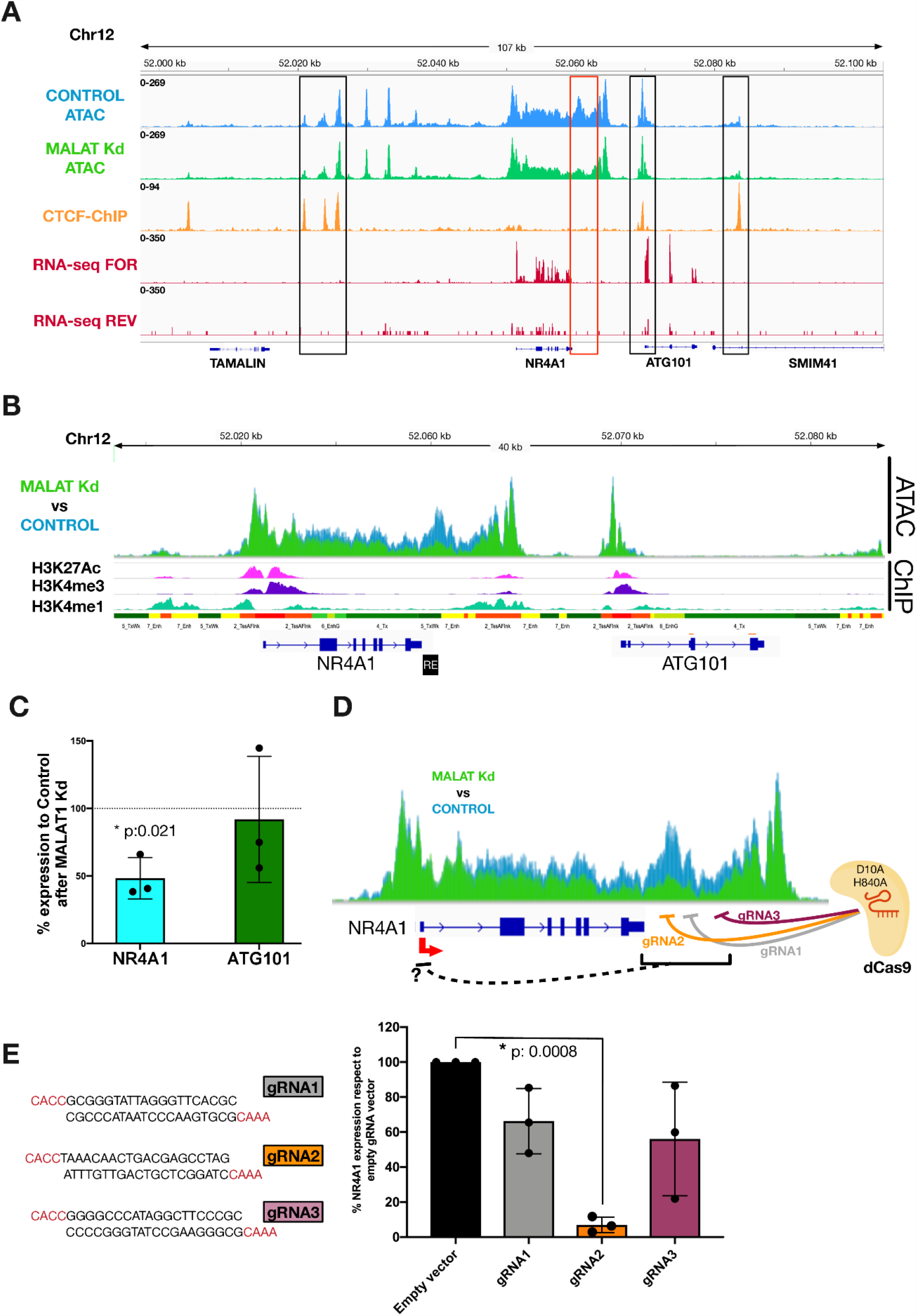
MALAT1 modulates the accessibility of a *NR4A1* downstream regulatory element. **A)** IGV screenshot of the *NR4A1* topologically associated domain defined by CTCF and ATAC peaks. **B)** Chromatin states over the NR4A1 gene. **C)** Real-time PCR expression analysis of NR4A1 and ATG101 after MALAT1 Kd. **D)** Scheme depicting the CRISPR-i approach using 3 gRNAs against the *NR4A1* downstream regulatory element. **E)** Guide RNAs sequences and real-time PCR quantification of NR4A1 expression after CRISPR-i assays. *Significant differences are shown with p-values obtained from t-tests analyses.

MALAT1 Kd resulted in a significant downregulation of the *NR4A1* gene, without changing the expression rates of *ATG101* (Figure 1K, Supplementary Table 2). This effect was verified by RT-qPCR (Figure 3C), suggesting a direct, MALAT1-dependent loss of chromatin accessibility at a novel NR4A1-downstream regulatory element (Figure 3B, black box). To test the functionality of the proposed RE, we used the CRISPR-i system, consisting of the catalytically inactive Cas9 (dCas9) and three independent guide RNAs (gRNAs) targeting the *NR4A1* downstream regulatory element (Figure 3D). Guide RNAs were designed to bind directly to the proposed RE (gRNA2), or 1 kbp (gRNA1) and 2 kbp (gRNA3) downstream of the RE. HeLa cells were independently transfected with the three guides RNA plasmids (Supplementary Figure 3A) and dCas9. Real-time PCR analysis showed repression of NR4A1 expression with all three guide RNAs, with gRNA 2 hitting the RE being the most efficient guide RNA (Figure 3E). The results indicate the presence of a potential novel downstream regulatory element, regulating the expression of the NR4A1 gene.

Next, we tested whether the MALAT1 RNA is physically associated with the downstream regulatory element of the NR4A1 gene, asking whether the lncRNA maintains DNA accessibility at this locus. Using the RAP assay we pulled down MALAT1 from HeLa nuclear extract (Supplementary Figure 2B). However, we did not find a direct-physical interaction of MALAT1 with the *NR4A1* downstream regulatory element (Supplementary Figure 2C), suggesting that additional factors mediate chromatin closure.

Taken together, the results show that MALAT1 regulates the accessibility of a novel downstream regulatory element, fine-tuning the expression of NR4A1 transcripts in HeLa cells.

### Chromatin accessibility of the *NR4A1* downstream regulatory element correlates with NR4A1 expression in different cancer types

To address the functional role of the newly identified downstream regulatory region of the NR4A1 gene we screened the TCGA cancer database. Indeed, we observed an overlap between the RE and 2 ATAC-seq peaks derived from TCGA data (58) (Figure 4A). The intensity of both ATAC-seq peaks does positively correlate with NR4A1 expression overall cancer types (Figure 4B). Furthermore, analyzing the accessibility of the two ATAC peaks in individual cancer types, showed that the accessibility of this region was cancer-type specific (Figure 4C). The highest level of accessibility was observed for Pheochromocytoma and Paraganglioma (PCPG) and the lowest ATAC-seq scores were found for Kidney Renal Papillary cell carcinoma (KIRP) (Figure 4C). These cancer-specific effects led us to evaluate the correlation of accessibility for both peaks versus NR4A1 expression in different cancer types. Here we identified a strong correlation for Breast Invasive Carcinoma (BRCA) when compared to other cancer types (Figure 4D). The direct correlations between NR4A1 expression, and the accessibility of *NR4A1* downstream regulatory element (NR4A1-RE), define a MALAT1/NR4A1 axis that, according to our initial results in HeLa cells and the bioinformatic analysis of TCGA data, suggest the MALAT1 regulation of NR4A1 expression through accessibility changes in the downstream element in a cancer-specific manner.

**Figure 4.**
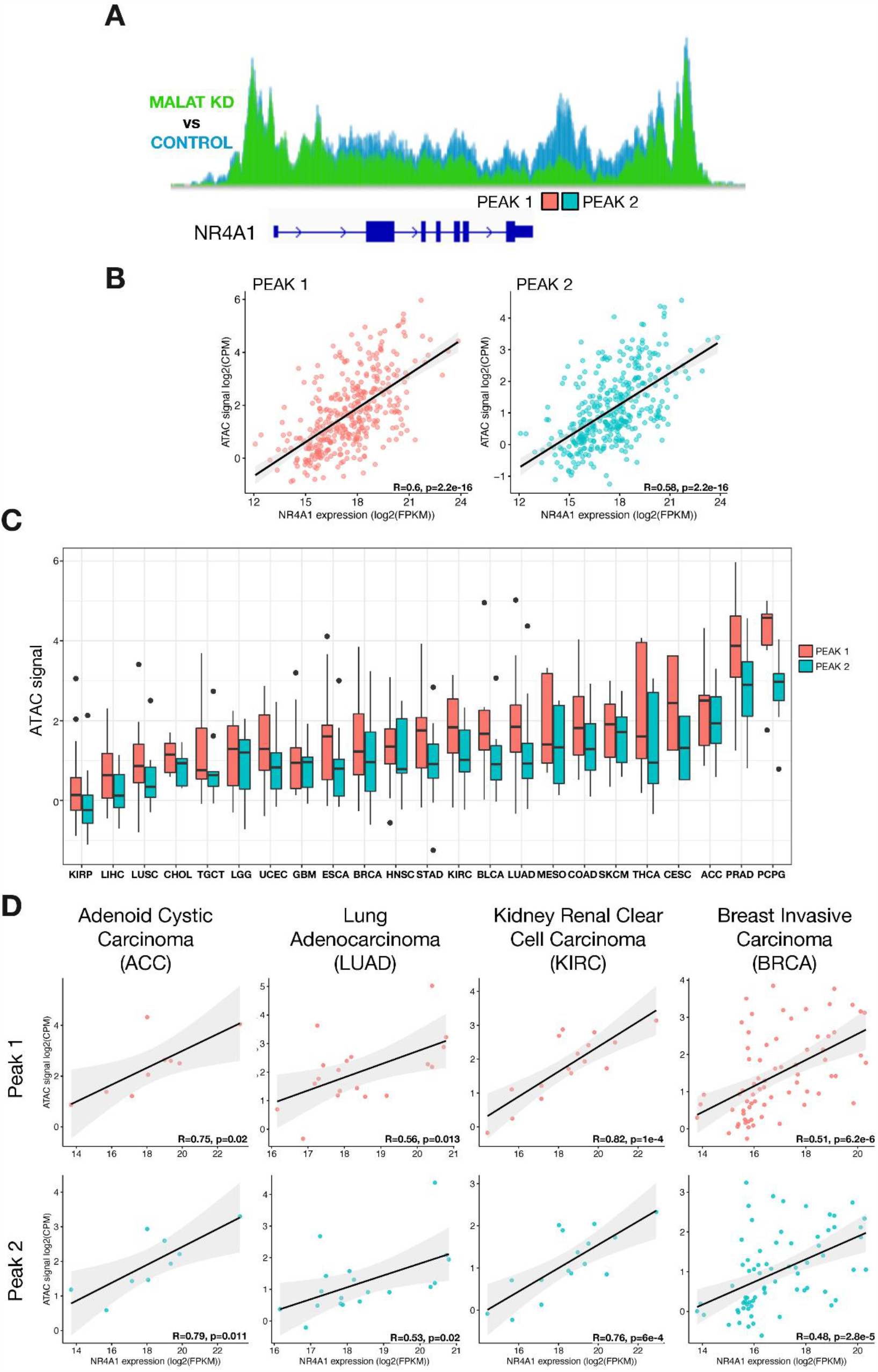
Association of the accessibility of the downstream regulatory element and NR4A1 expression in specific cancer types. **A)** Schematic representation of the TCGA-derived peaks found over the *NR4A1* downstream regulatory element. **B)** Correlation of the TCGA-derived peaks over the *NR4A1* downstream regulatory element versus NR4A1 expression on different TCGA cancer-types data. **C)** Accessibility quantification of the TCGA-derived peaks found over the *NR4A1* downstream regulatory element in specific cancer types. **D)** Correlations of the accessibility of TCGA-derived peaks over the *NR4A1* downstream regulatory element and NR4A1 expression on individual TCGA cancer types.

### The MALAT1-NR4A1 axis is functional on breast MCF7, but not on pancreatic PANC1, cancer cells

To prove the hypothesis of a cancer-type specific MALAT1/NR4A1-RE axis of NR4A1 expression, we evaluated the effect in specific cancer cell types. We tested cancer types exhibiting a high correlation of MALAT1 expression and NR4A1-RE accessibility, such as BRCA (Figure 4E), versus low- or no-correlation, like pancreatic adenocarcinoma (PDAC) (Figure 5A). To assess the functionality of the MALAT1/NR4A1-RE axis, we transfected Gapmers to downregulate MALAT1 in MCF7 breast cancer cells and PANC1 pancreatic adenocarcinoma cells (Figure 5B). MALAT1 Kd induces a significant decrease in NR4A1 expression in MCF7 cells, while no difference compared to the controls was observed in PANC1 cells (Figure 5C). Furthermore, we evaluated the regulatory function of the NR4A1 downstream regulatory element. As described in Figure 3C, we transfected dCas9 and gRNA2 into MCF7 cells (Figure 5D) and evaluated the expression of NR4A1. A significant reduction in the expression of NR4A1 was observed (Figure 5E). Meaning that the NR4A1 downstream element is sufficient to modulate the expression of the NR4A1 gene in MCF7 cells.

**Figure 5.**
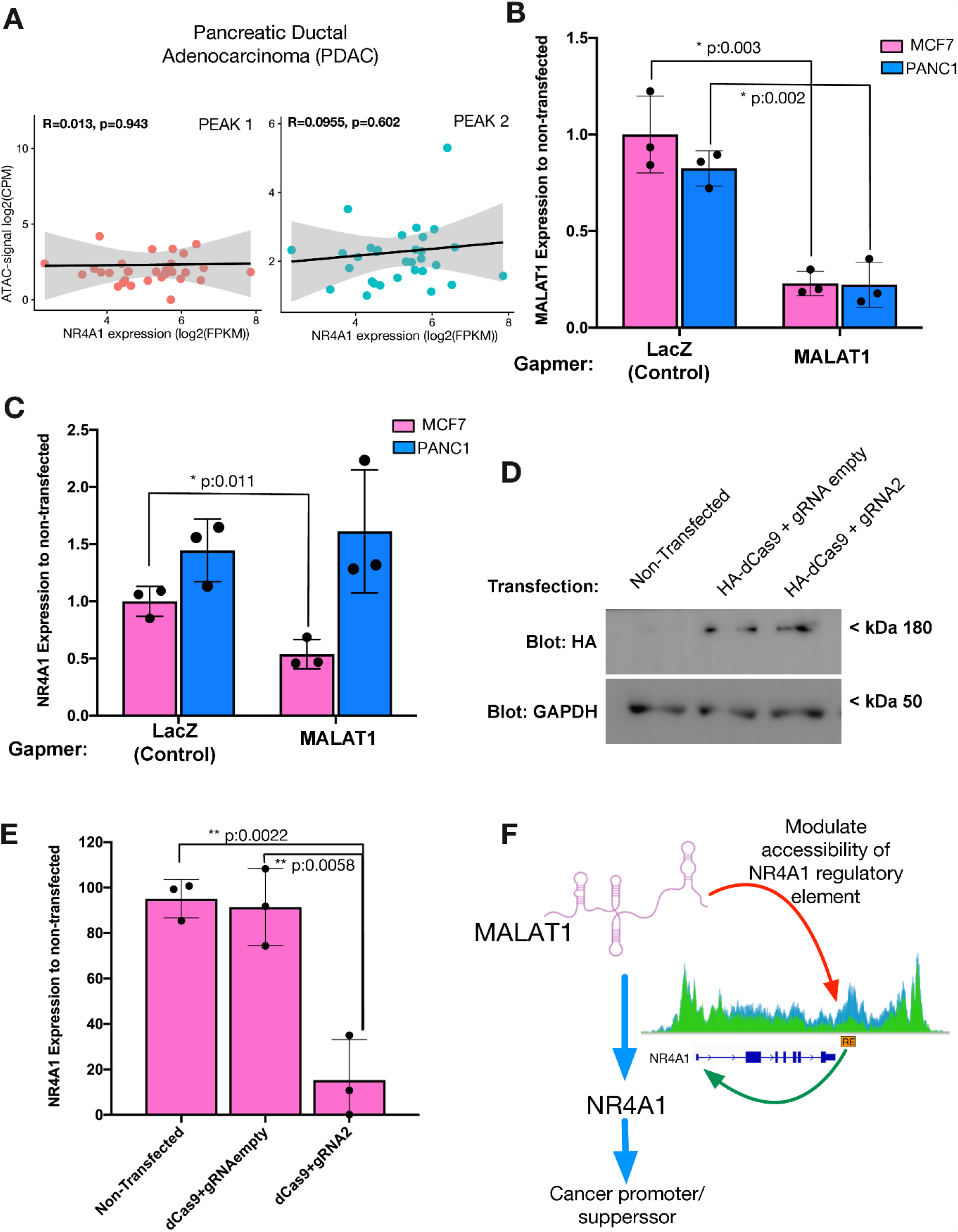
MALAT1 regulation of NR4A1 occurs on breast cancer cells, but not on pancreatic cells. **A)** Correlations of the accessibility of TCGA-derived peaks over the *NR4A1* downstream regulatory element versus NR4A1 expression on PDAC. **B)** MALAT1 downregulation on MCF7 and PANC1 cells using Gapmers. **C)** Real-time analysis of NR4A1 expression after MALAT1 Kd on MCF7 and PANC1 cells using Gapmers. **D)** Western blot analysis of the HA-tag expression to verify the dCas9 transfections on MCF7 cells. **E)** CRISPR-i examination of the gRNA2 over the *NR4A1* downstream regulatory element by real-time PCR. **F)** Scheme representing the molecular mechanism describing the MALAT1 modulation of NR4A1 through the accessibility of a downstream regulatory element.

## Discussion

Chromatin-associated lncRNAs are of key importance in the regulation of cellular transcriptional levels. During cancer onset and progression, altered transcriptional programs trigger uncontrolled intracellular signaling that normally induces cell de-differentiation and proliferation (36–38). In this context, MALAT1 has been involved in different cancer types, acting as a transcriptional regulator of oncogenes and tumor suppressors (6,17,18). Our work focused on deciphering the molecular mechanism of gene expression regulation of the chromatin-associated lncRNA MALAT1.

### MALAT1-dependent changes in chromatin accessibility

We found that the presence of MALAT1 was essential for the expression of many protein-coding genes. With the knock-down of the lncRNA alterations in chromatin accessibility of a large number of loci were observed. Such an increase or decrease in chromatin accessibility after a lncRNA Kd has been described before (34,35,39,40), confirming that lncRNAs are active chromatin regulators. Nevertheless, to our knowledge, there is no study showing the impact of MALAT1 on chromatin accessibility. The ATAC-seq data identified the largest changes in chromatin accessibility downstream of the *NR4A1* gene. The local decrease within an open 36 kbp region, was associated with the repression of the NR4A1 gene expression. We showed that this MALATA1-dependent ATAC-seq peak presents a new NR4A1 downstream regulatory element (NR4A1-RE), as CRISPR-i experiments targeting the NR4A1-RE also downregulated the expression of the gene. Similar types of REs have been reported to regulate *Cja1* (Cx43) and mouse *Globin* genes (42,43). MALAT1 RAP assays could not detect the physical interaction of MALAT1 with the NR4A1 locus, suggesting that MALAT1 does not directly maintain the open chromatin structure of the NR4A1-RE in HeLa cells. MALAT1 CHART-seq data did not identify the interaction of MALAT1 with the NR4A1-RE in ER-positive MCF7 breast cancer cells, confirming our findings (Supplementary Figure 4A) (16). Our ATAC-seq data showed that the NR4A1 promoter and the NR4A1-RE of NR4A1, correspond to a region with the highest level of chromatin accessibility of the whole chromosome 12. The decrease in accessibility of the NR4A1-RE after MALAT1 Kd involves additional factors that maintain this region accessible under normal conditions. Data retrieved from the ChIP-atlas (https://chip-atlas.org/) in HeLa cells, revealed that the transcription activator BRG1, the catalytic subunit of the SWI/SNIF chromatin remodeling complex, binds along the whole NR4A1 gene body, including the NR4A1-RE (Supplementary Figure 4B). A direct MALAT1-BRG1 interaction was previously shown to increase the inflammatory response in hepatocellular carcinoma (44). Besides BRG1, two other transcription factors involved in the super elongation complex of RNA polymerase II, AFF4 and ELL2 (45), are also binding to the NR4A1 gene body and the NR4A1-RE (Supplementary Figure 4C). MALAT1 Kd resulted in a minor decrease of BRG1 and AFF4 expression, but with a significant down-regulation of ELL2 (Supplementary Figure 4D and Supplementary Table 2). The putative MALAT1-BRG1 interaction and the slight expression changes could explain the MALAT1-dependent changes in chromatin accessibility at the NR4A1-RE. Nevertheless, apart from additional protein factors, we cannot rule out the possibility that other ncRNAs are involved in maintaining this region accessible.

A recent report describes the expression of an antisense lncRNA to NR4A1, NR4A1AS, in colorectal cancer cells (46). The promoter region of NR4A1AS1 coincides with the NR4A1-RE, making this lncRNA an interesting candidate to follow up in a subsequent study. However, our RNA-seq data did not reveal the presence of this lncRNA in HeLa cells. Moreover, the result was verified by real-time PCR assays in HeLa and also MCF7 cells, not detecting the NR4A1AS RNA (data not shown). These results indicate that NR4A1AS is not involved in the presented model systems, but most probably, the antisense RNA plays a tissue-specific role.

### Impact of the MALAT1-NR4A1 axis on breast cancer development

TCGA data revealed that the chromatin accessibility of the NR4A1-RE is directly correlated with *NR4A1* expression in several cancer types as we observed in HeLa cells. Here, we focused on breast carcinoma cells due to their high correlation. We were able to recapitulate the MALAT1-NR4A1 axis in ER-positive MCF7 breast cancer cells and did not find it in pancreatic adenocarcinoma PANC1 cells, as predicted by the TCGA data. The function of MALAT1 expression in breast cancer cells is still a matter of debate. Both, oncogenic and tumor suppressor roles have been described for this lncRNA (47). Despite this disagreement, MALAT1 was shown to be chromatin-associated in two different breast cancer types, ER-positive MCF7 and triple-negative breast cancer (TNBC) MDA-MB-231 cells (16,30). Our results agree with the role of MALAT1 as a tumor suppressor, by maintaining the NR4A1 expression in MCF7 cells. On the one hand, NR4A1 upholds the ERK signaling at low levels and preserves the levels of pro-apoptotic proteins, regulating proliferation and controlled cell death, respectively (27). Through these mechanisms, NR4A1 seems to retain the tamoxifen sensitivity of MCF7 cells (27). On the other hand, in TNBC MDA-MB-231 cells, NR4A1 acts in a different manner. NR4A1 is promoting tumor invasion and metastasis by activating transforming growth factor beta (TGF-R) signaling (26). These results suggest that depending on the cellular environment, the MALAT1-NR4A1 axis can work as a tumor promoter or suppressor (Figure 5F).

The transcription factor NR4A1 has been described as an immediate early gene (IEG), which corresponds to a group of genes that rapidly respond to stress through their fast and transient transcriptional stimulation (48). Replication stress in the absence of checkpoints and quality control machinery is one of the first steps leading to genomic instability during cancer development (49). Comparing non-malignant MCF10 breast epithelial cells with TNBC cells MDA-MB-231, and circulating patient-derived breast tumor cells, NR4A1 has been recently reported to modulate the IEG’s expression through the interaction with their gene bodies and suppressing their transcriptional elongation. While under oncogenic replication stress conditions, NR4A1 is released, triggering a burst of IEG’s expression (50).

Together, the data pinpoints to a mechanism where MALAT1 modulates the accessibility of a downstream regulatory element (NR4A1-RE) to determine NR4A1 expression levels. The presence of MALAT1 seems to fine-tune the NR4A1 expression levels, which in turn, results in low proliferative signaling and pro-apoptosis cues on ER-positive MCF7 cells. The discovery of the MALAT1-NR4A1 axis increases their potential as therapeutic targets for ER-positive breast cancer and drug-resistance treatments.

## Acknowledgments

This work was funded by the Agencia Nacional de Investigación y Desarrollo (ANID, FONDECYT de Iniciación 11190246 to R. M.), and by the Deutsche Forschungsgemeinschaft (SFB960 and LA1331 to G. L.). The results shown here are in part based upon data generated by the TCGA Research Network https://www.cancer.gov/tcga.

## Material and Methods

### Cell Culture

HeLa, MCF7, and PANC1 cells were cultured in HyClone Dubelco’s Modified Eagle Medium (DMEM), supplemented with glucose 4.5 g/L, L-Glutamine 584 mg/L, sodium pyruvate 110 mg./L, and 10% Fetal bovine Serum. Cells are maintained under constant conditions of 37°C and 5% CO_2_ and subcultured using Trypsin 0.25%.

### Nuclear preparations

HeLa nuclear preparations were performed following the REAP (Rapid, Efficient And Practical) protocol (51) with modifications for crosslinked cells. In brief, twenty million HeLa cells were harvested in 1 mL of PBS 0.2% NP-40 supplemented with protease and RNase inhibitors. Then, two washes of the cytoplasmic fraction are made by pipetting up/down, vortexing, 10 times pipetting with a 1 mL syringe and 23G needle, and 12.000xg centrifugation for 30 seconds. Finally, the nuclear pellet was resuspended in 500 μL of RIPA buffer (10 mM Tris-HCl [pH 7.6], 1 mM EDTA, 0.1% SDS, 0.1% Sodium Deoxycholate, 1% Triton X-100) for sonication.

### Western Blot

Protein extracts from HeLa and MCF7 cells were obtained after homogenization with RIPA buffer supplemented with protease inhibitors by pipetting and sonication at 4°C. The proteins were resolved by SDS-PAGE and transferred to PVDF membranes. Membranes were blocked with 5% milk in TBST (TBS with 0.1% Tween-20 ®), probed with antibodies against H3K27ac, Tubulin, and HA-tag, and then with horseradish peroxidase-coupled secondary antibodies. Luminescent blot signals were obtained by incubating with the peroxidase substrate and captured on a Syngene G:box Gel documentation system.

### Real-time PCR

Total RNA (1 μg) from HeLa, MCF7, and PANC1 cells, or 500 μg of RNA from HeLa ChRIP and input samples, were used for cDNA preparation with the RevertAid RT Kit (Thermo Scientific). Real-time PCR reactions were prepared in 0.1 mL tubes with 20 μl of the final volume containing specific primers (MALAT1, NR4A1, GAPDH), variable amounts of cDNA (depending on the target), 10 μl of the 2X JumpStart™ ReadyMix™, 1 mM MgCl_2_, SyberGreen (1:400.000 from the 10.000X stock, Thermo Scientific). Reactions are analyzed on a Rotor-Gene RG3000 thermal cycler.

### Transfections

HeLa cells were reverse-transfected with Lipofectamine 2000 for Gapmer-mediated MALAT1 downregulation and CRISPR-I assays. Gapmer transfections were carried out using 1.5 μl of 200 μM Gapmers against MALAT1 or LacZ and 5 μl of lipofectamine for 1.2 million cells on a 6-well plate, for 24 hours post-transfection assays. For 48 hours post-transfection experiments, we used half the amount of cells. For CRISPR-i, we used 5 μl of lipofectamine, 750 μg of dCas9 plasmid (#61355 Addgene), and 750 μg of each gRNA plasmid (#47108 Addgene) for 0.5 million cells on a 6-well plate. The cells were harvested 48 hours post-transfections.

MCF7 cells were plated and transfected with Lipofectamine 3000 for Gapmer-mediated MALAT1 downregulation and CRISPR-I assays. Gapmer transfections were carried out using 2 μl of 200 μM Gapmers, against MALAT1 or LacZ, 2 μl of lipofectamine, and 1 μl of P3000 reagent, for 0.5 million cells seeded the day before on a 6 well plate. Cells were harvested 24 hours post-transfection. For CRISPR-i, we used 2 μl of lipofectamine, 650 μg of dCas9 plasmid (#47106 Addgene), 650 μg of each gRNA plasmid (#47108 Addgene), and 2.5 μl of P3000 Reagent, for 0.5 million cells seeded the day before on a 6 well plate. The cells were harvested 48 hours post-transfections.

### Chromatin-RNA Immunoprecipitation (ChRIP)

Twenty million HeLa cells were crosslinked with 1% formaldehyde for 8 minutes at room temperature with gentle agitation. Then, the reaction is quenched with 125 mM of glycine. Nuclei samples were prepared and sheared on a Bioruptor sonicator (Diagenode) at a medium intensity to obtain chromatin sizes between 200-800 bp. Sonicated samples were centrifuged at 8.000 x g for 30 seconds to eliminate nuclei debris, and the supernatant was used for pre-clear and immunoprecipitation. Pre-clearing was performed by incubating the sonicated samples with protein-G dynabeads ® (Thermofisher) for 1 hour at 4°C on a rotator wheel. Pre-cleared samples were incubated with 5 μg of the H3K27Ac (Ab4729 Abcam), and as a negative control, we used the same H3K27Ac antibody preabsorbed with the peptide to raise it (Ab24404 Abcam). Immunoprecipitations were incubated for 4 hours at room temperature on a rotator wheel. Five percent of the immunoprecipitated samples were used for western blot analysis, and the rest of the sample was used for RNA purification.

### Custom genome

A custom version of the hg38 human genome was used for all sequencing analysis, where all repeats of ribosomal genes were masked with N bases and a single repeat of the ribosomal gene was kept as an extra chromosome named “ChrR” (59).

### Annotation

A custom version of the gencode v27 Homo Sapiens annotation was used, with ribosomal RNA present one time on an extra chromosome - ChrR, to fit our custom genome.

### ChRIP-sequencing and pulldown analysis

Raw files were first checked for quality and adapter contamination with fastqc v0.11.9 (60) and multiqc v1.13 (61). Following the quality check, reads were aligned to the reference genome, hg38, using Bowtie2 (62) and the following flags: --local --very-sensitive-local. Unmapped reads and reads with lower than 20 MAPQ mapping quality and reads mapping to ribosomal RNA were removed for downstream analysis, using Deeptools alignmentSieve v3.1.2 (63). Next, read counts were obtained with FeatureCount from the Subread package v2.0.0 (64), using the following flags: -s 1 -t gene -g gene_id. Differential binding analysis between the knockdown samples and the controls was performed in R v4.1.2 with the DEseq2 R package (65). The cutoff to determine significantly enriched transcripts was Log fold change >0.5 and FDR <0.1.

### RNA purification

Total RNA purification for cDNA preparations was performed following TRIzol ® Reagent (Ambion by Life Technologies) manufacturer instructions. For ChRIP assays, RNA purification from immunoprecipitated samples was extracted by resuspending the beads on proteinase K buffer (10% SDS, 50 mM EDTA, 10 mM Tris-Cl pH 7.4), and incubating them with 50 μg of Proteinase K for 1 hour at 50°C with gentle agitation. Then, the crosslinking was reversed at 65°C for 2 hours, followed by RNA purification with TRIzol and Isopropanol precipitation.

### RNA-sequencing and transcriptome analysis

Raw files were first checked for quality and adapter contamination with fastqc v0.11.9 (60) and multiqc v1.13 (61). Following the quality check, adapters were removed using Trimmomatic v0.38 (66), with the following flags: PE -phred33 ILLUMINACLIP:TruSeq3-PE-2.fa:2:30:10 LEADING:3 TRAILING:3 MINLEN:30 SLIDINGWINDOW:4:20 and cutadapt v4.0 (67) for library- and sequencer-specific trimming, with the following flags: -u 5 -a “A{100}” -a “G{100}” -a “T{100}”. Reads were then aligned to the reference genome, hg38, using STAR v2.7.2a (68) and the following flags: --sjdbGTFfile gencode.v27.annotation.gtf --outFilterType BySJout --outFilterMultimapNmax 20 --alignSJoverhangMin 8 --alignSJDBoverhangMin 1 –outFilterMismatchNoverReadLmax 0.04 --alignIntronMin 20 --alignIntronMax 1000000 --alignMatesGapMax 1000000 -- outSAMtype BAM SortedByCoordinate --outFilterScoreMinOverLread 0 -- outFilterMatchNminOverLread 0 --outFilterMatchNmin 0. Unmapped reads and reads with lower than 30 MAPQ mapping quality were removed for downstream analysis, using samtools v1.90 (69). Next, read counts were obtained with FeatureCount from the Subread package v2.0.0 (64), using the following flags: -p -s 1 -t exon -g gene_id -C -B. Differential expression analysis between the knockdown samples and the controls was performed in R v4.1.2 with the DESeq2 R package (65), with apeglm shrinkage estimation for ranking of genes and visualization (70). Differential expressed genes were filtered with absolute Log fold change >1.0 and FDR <0.1.

### ATAC-seq assays, sequencing and analysis

ATAC assay was performed following the instructions of the Omni-ATAC protocol (52), using the Nextera DNA Flex Library prep kit (Illumina). Tagmented DNA was analyzed on a bioanalyzer and sequenced on a Next-Seq 550 device in the Genomics core unit of the Regensburger Centrum für Interventionelle Immunologie.

Raw files were first checked for quality and adapter contamination with fastqc v0.11.9 (60) and multiqc v1.13 (61). Following the quality check, adapters were removed using Trimmomatic v0.38 (66). Paired-end reads were than mapped to the reference genome hg38, using Bowtie2 (62) with the following flags: --local --very-sensitive-local -X 5000 -- no-discordant –dovetail. Unmapped reads and reads with lower than 30 MAPQ mapping quality were removed for downstream analysis, using samtools v1.90 (69). Reads mapping to the chrM, blacklisted regions and PCR duplicates were removed using samtools and Picard, respectively. Finally, reads were shifted based on Tn5 cuts, using DeepTools v3.1.2 (63) alignmentSieve --ATACshift. Peak calling was performed using MACS2 v2.2.71 (71), with the following flags: callpeak -f BAMPE -g hs –broad –keep-dup all –buffer-size=100. Differentially accessible regions were obtained with DiffBind R package. Differentially accessible regions were filtered with absolute Log fold change >1.0 and FDR <0.01. Regulatory regions were assigned to genes, using the GREAT tool (72), by basal plus extension method (5 kb upstream, 1 kb downstream plus distal up to 50 kb).

### Correlations with TCGA data and pancreatic cancer data

The normalized ATAC-seq peak signals (ID: Peak1 = PRAD_72732 and Peak2 = PRAD_72733) of TCGA tumor samples were obtained from the ATAC-seq Hub (https://atacseq.xenahubs.net) (58). Corresponding gene expression quantification of matching donors was obtained using the TCGAbiolinks Bioconductor package (73). Here we used the upper quartile normalized FPKM values.

The accessibility signals for Peak1 and Peak2 were then plotted against the NR4A1 expression. For the PANC1 cell line, ATAC-seq and RNA-seq data were obtained through the GEO database (GSE124229 and GSE124231). First, the gene expression and accessibility signals were normalized to the library size for each patient. The accessibility signal for Peak1 and Peak2 was then plotted against the NR4A1 expression. On both, linear regression was used to model the correlation between the accessibility and gene expression signal in R v.4.1.2. Heatmaps were made using DeepTools v.3.4.3 (63) and pheatmap R package.

