## Supplemental figures for "The long non-coding RNA MALAT1 modulates NR4A1 expression through a downstream regulatory element in specific cancer-cell-types"

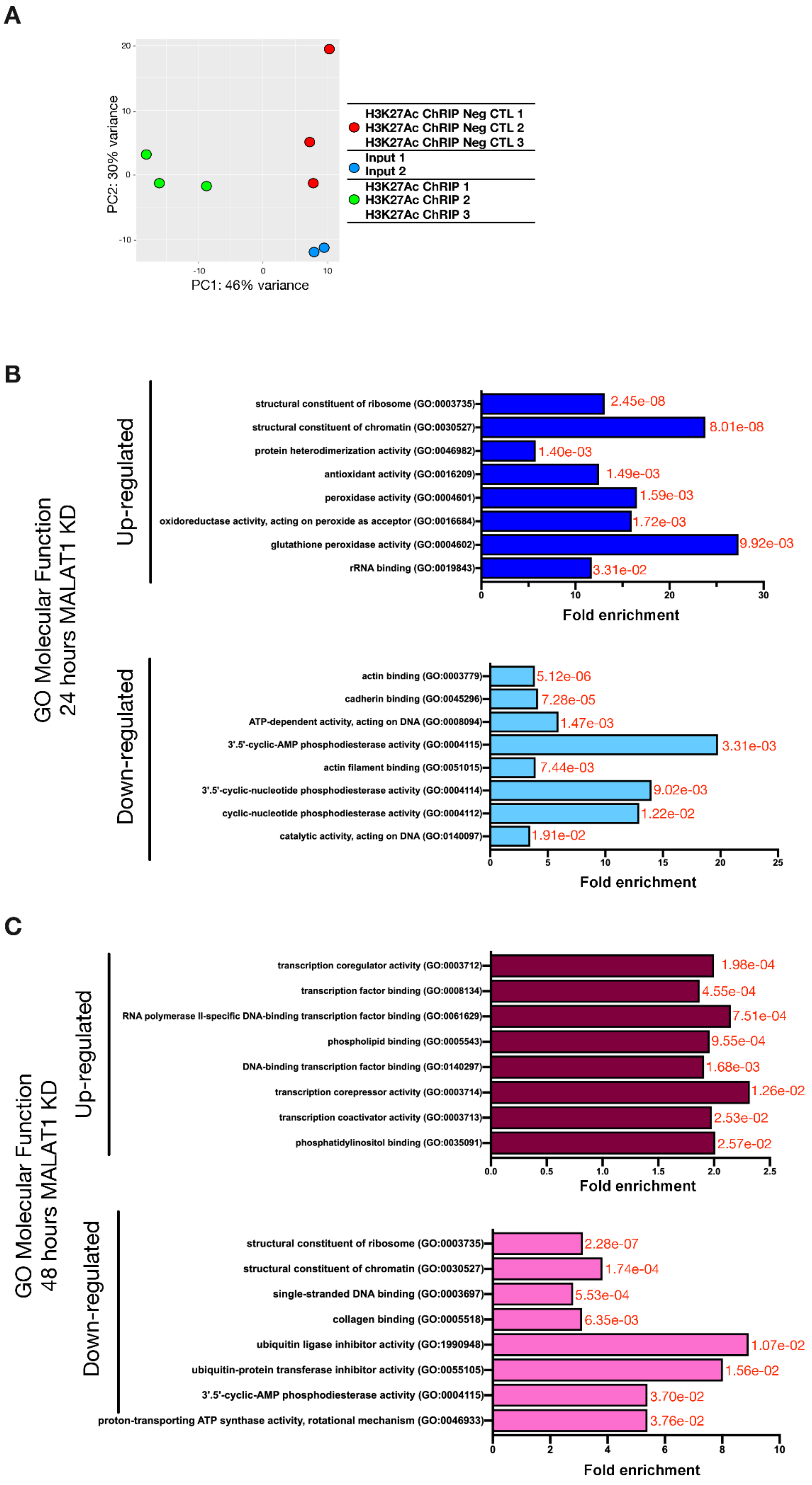


**Supplementary Figure 1. A)** Principal component analysis of the sequenced ChRIP samples. **B)** and **C)** Gene ontology analysis by overrepresentation test of transcripts up- and down-regulated after 24 and 48 hours MALAT1 Kd, respectively.

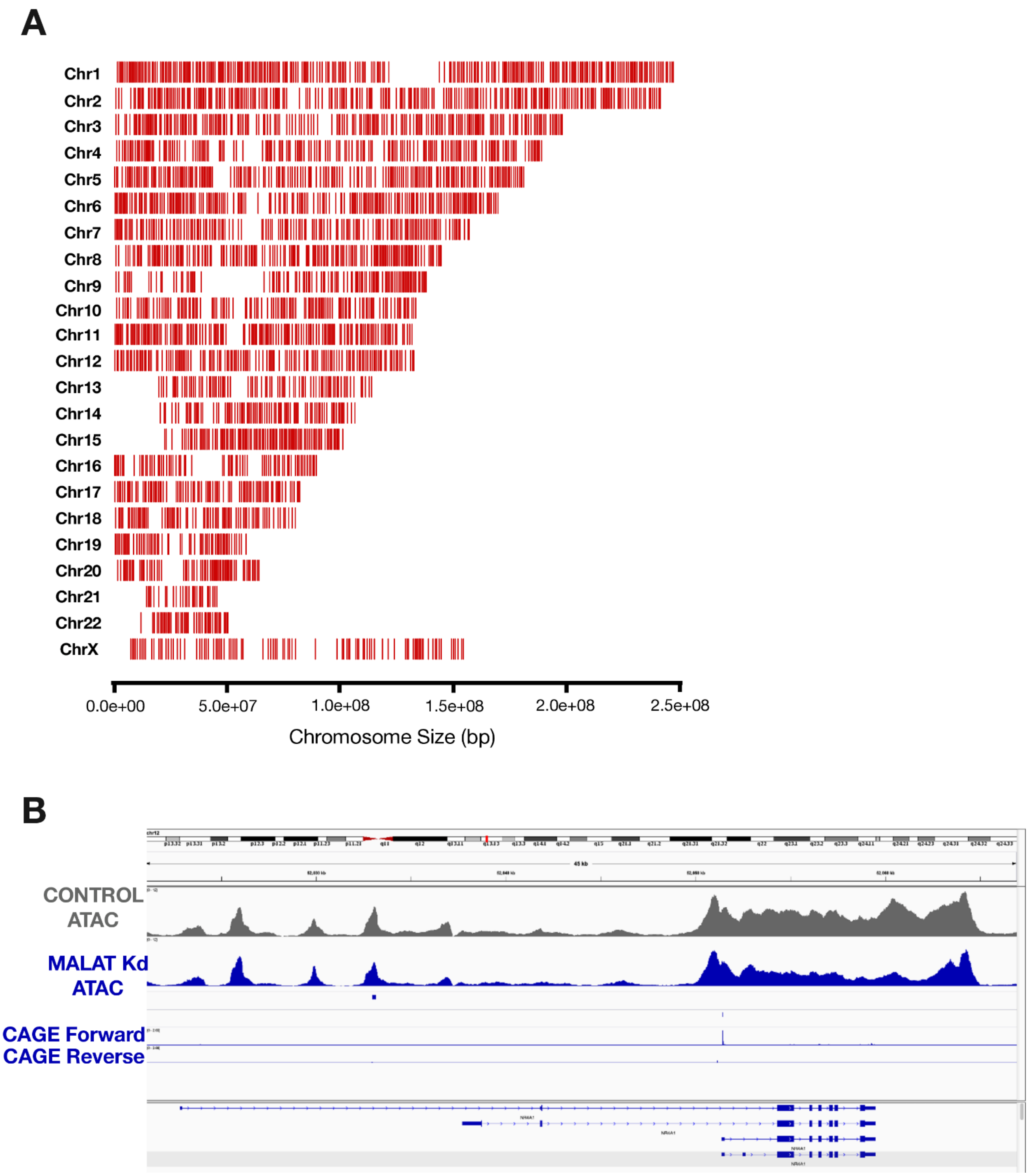


**Supplementary Figure 2. A)** Distribution of significant ATAC-seq peaks per chromosome. **B)** IGV screenshot depicting CAGE data to identify the NR4A1 transcriptional start site.

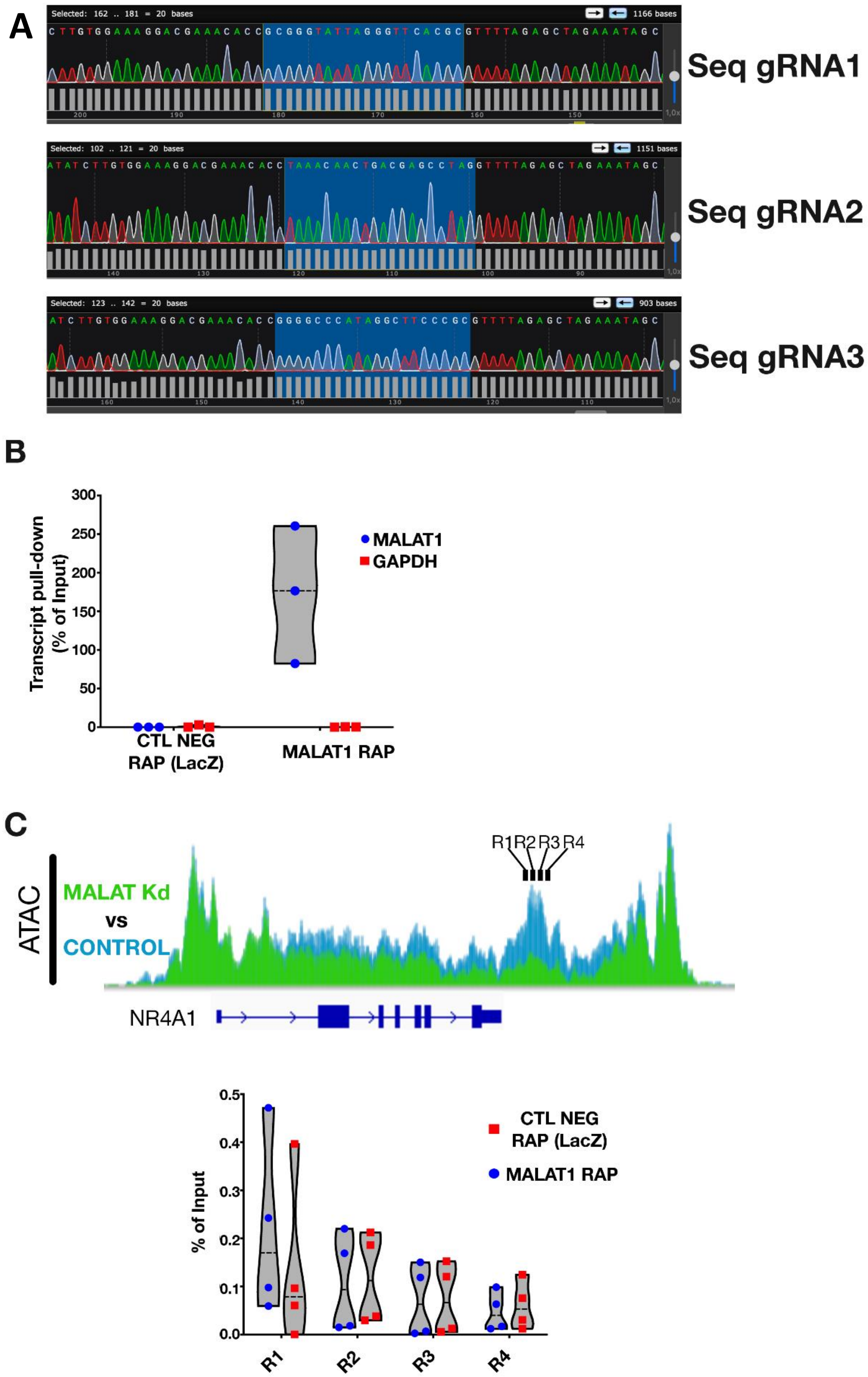


**Supplementary Figure 3. A)** Sequencing results for the plasmids coding for the gRNAs used in this study. **B)** RNA-affinity purification results for MALAT1 pull-down. Oligos against LacZ were used as a pull-down negative control, while GAPDH detection was performed for the detection of the negative control. **C)** Real-time PCR detecting 4 regions (R1 to R4) of the NR4A1 downstream regulatory element after MALAT 1 RNA-affinity purification.

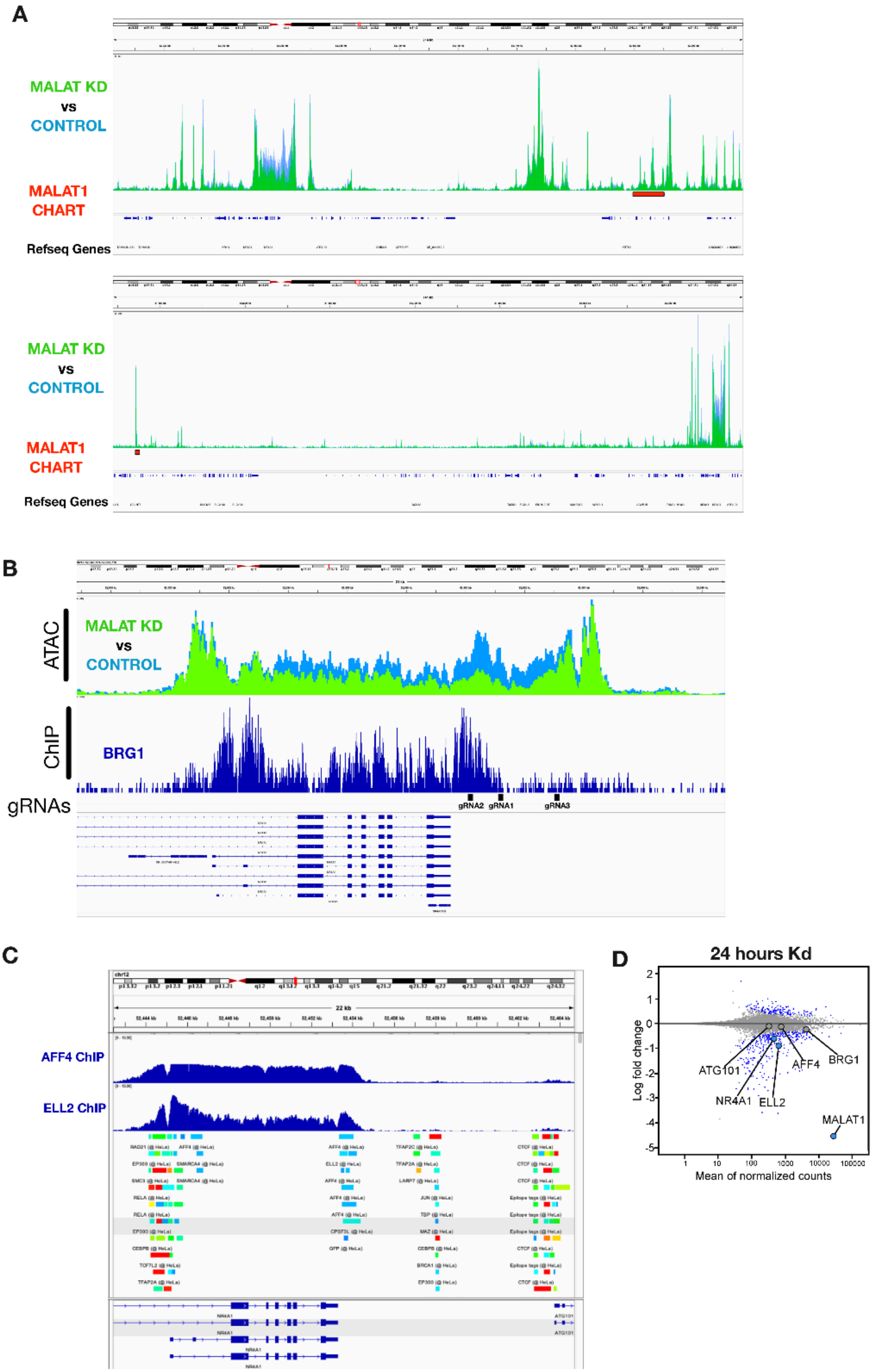
A
MALAT KD
vs
CONTROL
MALAT1
CHART
Refseq Genes
MALAT KD
vs
CONTROL
MALAT1
CHART
Refseq Genes
B
ATAC
MALAT KD
vs
CONTROL
ChIP
BRG1
gRNAs
gRNA2
gRNA1
gRNA3
C
AFF4 ChIP
ELL2 ChIP
22 kb
D
24 hours Kd
Log fold change
Mean of normalized counts
ATG101
AFF4
BRG1
NR4A1
ELL2
MALAT1

**Supplementary Figure 4. A)** MALAT1 CHART peaks on MCF7 cells downstream (up) and upstream (down) the *NR4A1* gene. **B)** BRG1 ChIP peaks over the *NR4A1* gene on HeLa cells. The specific targeting locations of the gRNAs for CRISPRi assays are indicated. **C)** AFF4 and ELL2 ChIP peaks over the *NR4A1* gene on HeLa cells. **D)** MALAT1 RNA-seq counts after 24 hours of MALAT1 Kd using specific Gapmers. The counts for BRG1, ELL2 and AFF4 are highlighted.
